# Drug design using unique conformations to preferentially target a specific site on collagen-bound MMP1

**DOI:** 10.64898/2026.05.14.725194

**Authors:** Anthony Nash, Chase Harms, Susanta K. Sarkar

## Abstract

Precise site-specific drug design remains a challenge in structure-based drug discovery. Most existing approaches screen for ligands to target binding pockets on a protein surface based on static structures obtained from techniques such as X-ray, NMR, cryo-EM, and AlphaFold. However, the structure-function paradigm is, in reality, a structure-dynamics-function relationship that determines a protein’s binding and activity. As such, drug screening or design without evaluating binding competition across the protein surface or considering the receptor’s dynamic, substrate-dependent conformational states is incomplete. Substrate-specific unique protein conformations are underexplored and offer novel opportunities for selective therapeutic targeting, though systematic workflows for identifying and exploiting such sites remain limited. Previously, we showed that collagen alters matrix metalloprotease-1 (MMP1) dynamics and that R405 is an allosteric residue on the MMP1 surface that exhibits strong dynamic correlations with its active site. Here, we present a substrate-specific allosteric drug-design framework that targets specific sites on a protein, using collagen-bound MMP1 as a model system. We determined the conformational dynamics of free and collagen-bound MMP1 using all-atom molecular dynamics (MD) simulations and categorized conformations into clusters of similar conformations. We then compared and identified unique conformations that occur only in collagen-bound MMP1 to design drugs against them using a machine-learning approach. The top three unique clusters were used to generate approximately 150,000 candidate compounds that were then screened against both the R405-centered region and all detectable binding pockets across the MMP1 surface. We have found several compounds that bind preferentially around R405 by at least 0.3 kcal/mol relative to competing sites across the surface. This strategy establishes a generalizable framework for designing ligands that preferentially target substrate-specific allosteric sites, providing new opportunities for precision therapeutics that modulate proteins in their biologically relevant functional states.

**Simple Summary:** In this paper, we establish a substrate-specific allosteric drug-design strategy that integrates all-atom molecular dynamics simulations, conformational clustering, machine-learning-based ligand design, and surface-wide binding-selectivity screening, using collagen-bound MMP1 as a model system. We show that collagen binding reshapes the conformational ensemble of MMP1, creating unique conformational states that are absent or inaccessible in the free enzyme. By identifying these substrate-specific conformations, generating ligands based on the corresponding dynamic fingerprints around the collagen-specific allosteric residue R405, and screening compounds across all binding pockets on the MMP1 surface, we demonstrate preferential targeting of the collagen-specific site relative to competing pockets. These results establish a generalizable framework for designing ligands that selectively recognize biologically relevant substrate-bound conformations rather than static protein structures alone. Substrate-specific allosteric targeting may enable selective modulation of individual protein functions while minimizing off-target interactions, providing new opportunities for precision therapeutics against dynamic protein systems.

## Introduction

MMPs are zinc-dependent enzymes involved in diverse functions in the human body [1]. MMPs pose a challenge for drug targeting because inhibiting the active site affects both intended and other critical functions. As such, early inhibitors targeting the catalytic zinc-binding sites lacked specificity and produced dose-limiting toxicities, motivating a shift toward exosites and allosteric regulatory regions [2]. Therefore, controlling one specific function of MMPs is necessary to reduce the toxicity of drugs that target MMPs. In this paper, we focus on MMP1, a collagenase that cleaves triple-helical collagen and plays a central role in tissue remodeling and pathological invasion. Structural studies of collagen-bound MMP1 revealed that both the catalytic and hemopexin domains contribute to its function [3], and allosteric communications from the hemopexin domain are essential for cleaving triple-helical type-1 collagen [4].

Previously, we showed that the presence of a bound substrate alters MMP1 dynamics and that there are substrate-specific allosteric residues distant from the active site, having strong correlations with the active site [5, 6]. Identification of the substrate-specific allosteric residues provided an opportunity to alter one MMP1 function without affecting its other functions. Note that the allosteric residues of MMP1 are different depending on what substrate is bound, including no substrate (free MMP1). For collagen-bound MMP1, we identified R405 as an allosteric residue that shows a strong dynamic correlation with the active-site residue E219. From a biophysical perspective, these observations indicate that substrate binding reshapes the underlying conformational ensemble of MMP1 and establishes long-range dynamic coupling between distal residues. Such ensemble redistribution is a hallmark of allosteric regulation in flexible multidomain proteins [7]. In a previous paper, we screened a subset of molecules from the ZINC database against the region around R405 on MMP1’s surface and showed that ligand-binding affinity at R405 depends on the presence of collagen, and that some molecules preferentially bind at the allosteric site rather than other binding pockets [8]. As such, in this work, we leveraged machine-learning-based drug design against R405 to design drugs that preferentially bind to the targeted site rather than other sites on MMP1.

Since the emergence of computational drug discovery in the 1970s, the field has seen remarkable progress enabled by molecular dynamics (MD) simulations [9], high-performance computing facilities [10], on-demand virtual libraries of drug-like small molecules containing billions of ligands [11], an increasing number of available crystal structures [12], structural predictions by AlphaFold [13], and recent rapid progress in artificial intelligence [14]. Advances in MD simulations have been particularly important for capturing time-dependent fluctuations in protein structures and resolving transient conformational states that are not accessible through static structural methods alone [15-17]. As with the uniqueness of human fingerprints [18], we identified conformational fingerprints specific to collagen-bound MMP1. These fingerprints represent physically distinct local conformational states within the broader ensemble and provide a quantitative description of how substrate engagement modifies accessible MMP1 conformations. Then, we used a machine-learning approach to generate ligands whose drug-like properties are based on the dynamic fingerprint unique to collagen-bound MMP1 at R405 [18]. We identified ligands that bind to R405 more strongly than to any other site on MMP1.

This work establishes a general framework for substrate-specific allosteric drug design that integrates the identification of substrate-specific allosteric sites, dynamic conformational fingerprinting at these sites, and drug design against the dynamic conformations around them. By ensuring preferential binding to unique conformations, this strategy addresses a major limitation of traditional structure-based drug design. The approach provides a quantitative, transferable method for selectively modulating individual protein functions while minimizing off-target interactions, and offers a pathway for designing precision therapeutics targeting substrate-dependent protein states.

## Results and discussion

### Collagen-bound MMP1 exhibits restricted conformational dynamics compared to free MMP1

To investigate changes in MMP1 dynamics due to collagen binding, we compared MD simulations of free and collagen-bound MMP1. Starting from the crystal structure (PDB ID: 4AUO) of collagen-bound inactive MMP1 [3], we mutated the residue A219 to E219 in order to simulate the dynamics of active MMP1, and constructed one model for free MMP1 and another for collagen-bound MMP1. Both models were equilibrated and simulated using three sets of randomly chosen starting velocities. Each production trajectory ran for 500 ns, yielding a cumulative simulation time of 1.5 μs for each system (see Methods for details).

The catalytic domain exhibited greater intrinsic mobility than the hemopexin domain in the free systems, consistent with prior experimental and computational observations. In collagen-bound systems, global domain fluctuations were significantly reduced, indicating that substrate interactions stabilize specific conformational states. This effect is supported by residue-level root-mean-squared fluctuation (RMSF) analysis, which shows decreased flexibility across multiple regions in the collagen-bound systems relative to the free systems. Residue flexibility was mapped onto the structures using RMSF values calculated every 20 frames. System structures were captured throughout the simulation trajectory and superimposed to provide a visual reference for the extent of flexibility across all residues (**Figure 1**).

**Figure 1.**
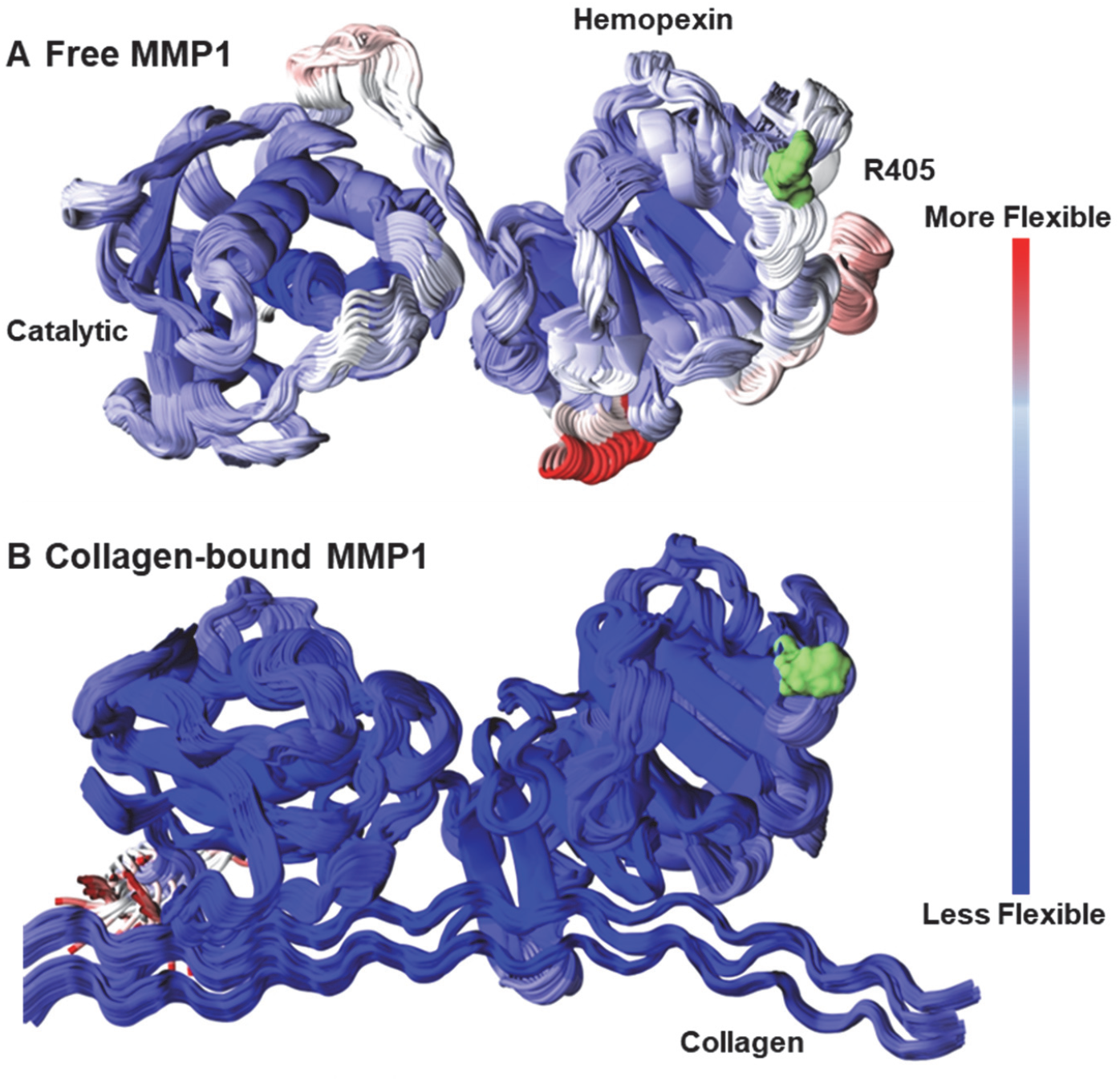
Dynamic flexibility of free and collagen-bound MMP1. Residue flexibility of **(A)** free MMP1 and **(B)** collagen-bound MMP1. The catalytic domain (F100–Y260) is on the left, the hemopexin domain (D279–C466) is on the right, and the two domains are connected by the linker region (G261–C278). Residue flexibility is mapped onto the structures using RMSF values calculated every 20 frames, with blue indicating the least mobile residues. Structures show the superposition of frames throughout the MD simulation. Collagen binding significantly reduces overall MMP1 flexibility. Residue R405 (in green) is a collagen-specific allosteric residue having strong correlations with the active site residue E219. We use the unique collagen-specific conformation around R405 to design drugs to preferentially bind at sites around R405.

These results demonstrate that collagen binding reshapes the dynamics landscape of MMP1 by restricting conformational variability and stabilizing catalytically relevant interdomain configurations. This shift in flexibility and the emergence of substrate-specific conformational states provide a mechanistic basis for identifying collagen-dependent allosteric regions suitable for substrate-selective ligand targeting. Collectively, these findings support the hypothesis that substrate binding generates unique structural states that can be exploited for substrate-specific allosteric drug design.

### Collagen-bound MMP1 has unique conformations distinct from those of free MMP1

To quantify substrate-dependent structural divergence, we compared representative conformations from clustered MD simulation frames of free and collagen-bound MMP1. **Figure 2A** presents a map of structural similarity scores computed between clusters derived from the dynamics of free and collagen-bound MMP1. Similarity measurements of conformation clusters were performed using conformations around the collagen-specific “fingerprint” of R405, within a sphere of radius 12 Å centered on its Cα atom. Structural similarity scores were calculated using Deeply Tough, a deep learning-based method for quantifying pocket and local structural similarity across protein conformations [19]. Scores ranged from −1 to 0, indicating maximal structural dissimilarity and identical local structural environments, respectively. **Figure 2B** shows the total similarity score across all free MMP1 clusters for each cluster of collagen-bound MMP1 conformations.

**Figure 2.**
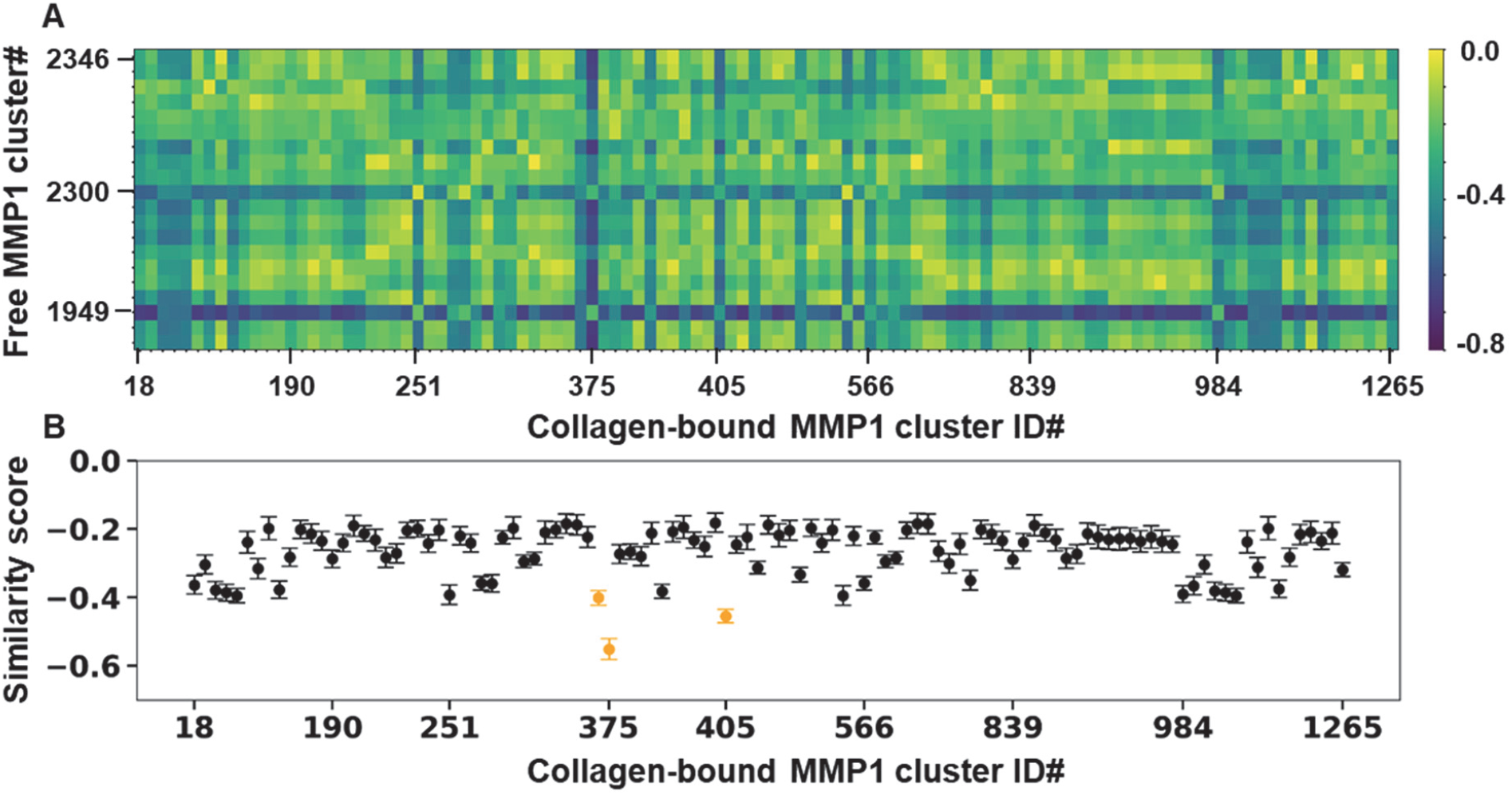
Identification of unique conformations of collagen-bound MMP1. (**A**) Pairwise similarity map between conformation clusters of free and collagen-bound MMP1. (**B**) The mean and standard error of the mean of similarity scores between each collagen-bound MMP1 conformation cluster and all clusters of free MMP1. The three clusters for collagen-bound MMP1 are shown in yellow.

We identified three conformational clusters with the lowest mean similarity scores relative to free-state conformations: -0.40 ± 0.02 (mean ± SEM, ID# 370), -0.55 ± 0.03 (mean ± SEM, ID# 375), and -0.46 ± 0.02 (mean ± SEM, ID# 405). These clusters, therefore, represent structural states uniquely enriched in the collagen-bound system and minimally sampled by free MMP1. We expected to generate distinct local geometries and physicochemical environments that influence ligand recognition. Consequently, structural features governing substrate-dependent drug binding are most likely to be captured within these uniquely collagen-associated conformational clusters, making them prime candidates for targeted ligand design.

### Collagen-specific allosteric fingerprints on MMP1 are dynamic

Traditional structure-based drug discovery commonly relies on screening compounds against static protein structures, implicitly assuming that binding pockets remain structurally stable and uniquely targetable across functional states [20]. However, proteins are dynamic and adopt an ensemble of conformations, and ligand or substrate binding can alter the distribution of conformations. The top three unique clusters (IDs 370, 375, and 405) in **Figure 2** differ in their three-dimensional arrangements, as shown in **Figure 3**. As such, collagen-specific allosteric fingerprints that surround defined residues represent dynamic, substrate-dependent screening targets. By incorporating conformational variability around a selected site, we aim to design ligands that preferentially bind to these biologically meaningful regions rather than nonspecifically interacting with multiple surface pockets. Such dynamic, site-selective targeting strategies may enable greater specificity and reduced off-target effects in precision therapeutic development.

**Figure 3.**
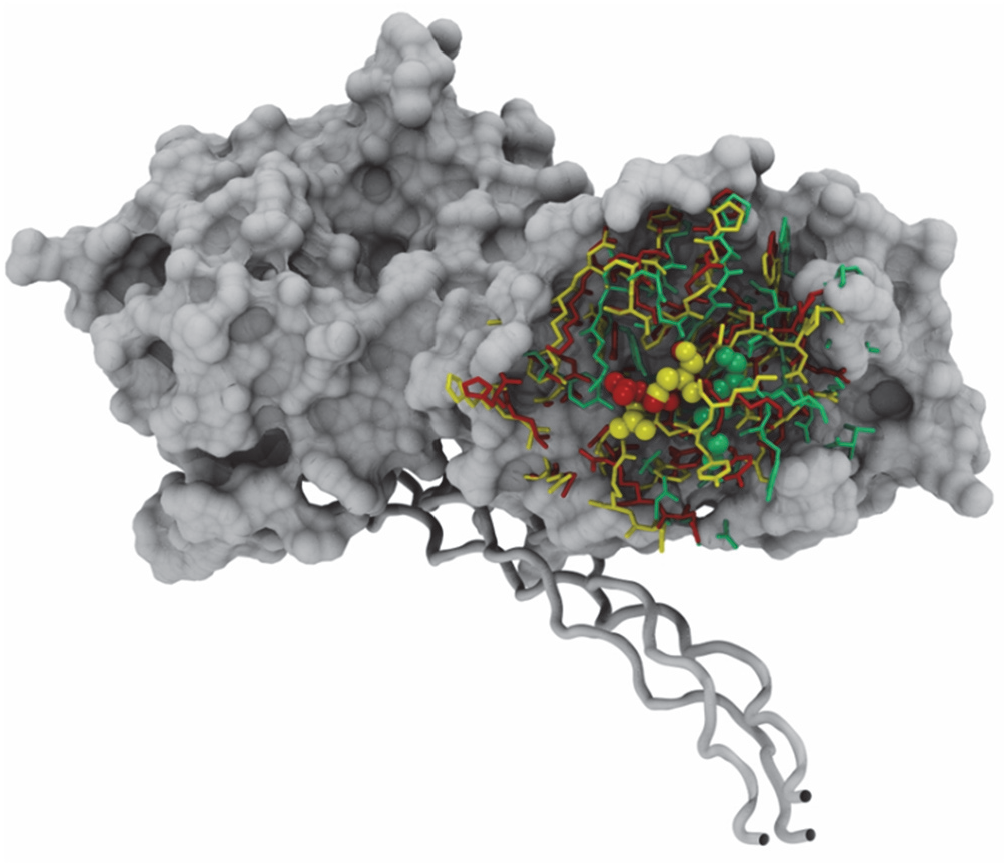
Dynamic collagen-specific allosteric fingerprints on MMP1. Three collagen-specific unique fingerprints are projected onto the protein surface, with each color representing one of three structurally distinct states. The position of R405 is indicated by spheres.

### Designed drugs using dynamic allosteric fingerprints yield better binding scores

For each of the three dynamic fingerprints in **Figure 3**, we designed ligands using an approach called Geometry-Complete Diffusion Model [18]. For each fingerprint, residues within 12 Å of R405 were defined as the local protein environment, and this region was used as a constraint for generating drug-like molecules. An iterative design and curation process was applied with a target of 50,000 unique ligands per fingerprint, while enforcing uniqueness both within each fingerprint-specific set and across all three sets. Molecules were subsequently filtered using RDKit-based criteria [21] and PoseBusters validation [22], with a threshold of 0.80 applied to retain ligands satisfying at least 80% of drug-likeness checks. Molecules failing these criteria were excluded, while accepted ligands were accumulated into fingerprint-specific datasets, yielding three structurally conditioned and chemically distinct ligand libraries.

The resulting ligand sets were evaluated with AutoDock GNINA docking [23], with five poses generated per ligand and centering the search space within 12 Å centered on R405 to fully encapsulate the fingerprint-defined region. This redocking framework enabled direct comparison of binding behavior across all generated compounds. The distribution of minimized affinity (kcal/mol) scores demonstrates that ligands generated from each fingerprint exhibit favorable binding characteristics within the targeted region (**Figure 4**). A substantial proportion of ligands fall within the range of approximately -2 to -5 kcal/mol, with a subset showing even more favorable values. As shown in **Figure 4**, the three fingerprint-derived ligand sets follow closely aligned score distributions, indicating that conditioning ligand generation on dynamic allosteric fingerprints yields compounds that consistently interact with the same local environment and comparable binding energetics.

**Figure 4.**
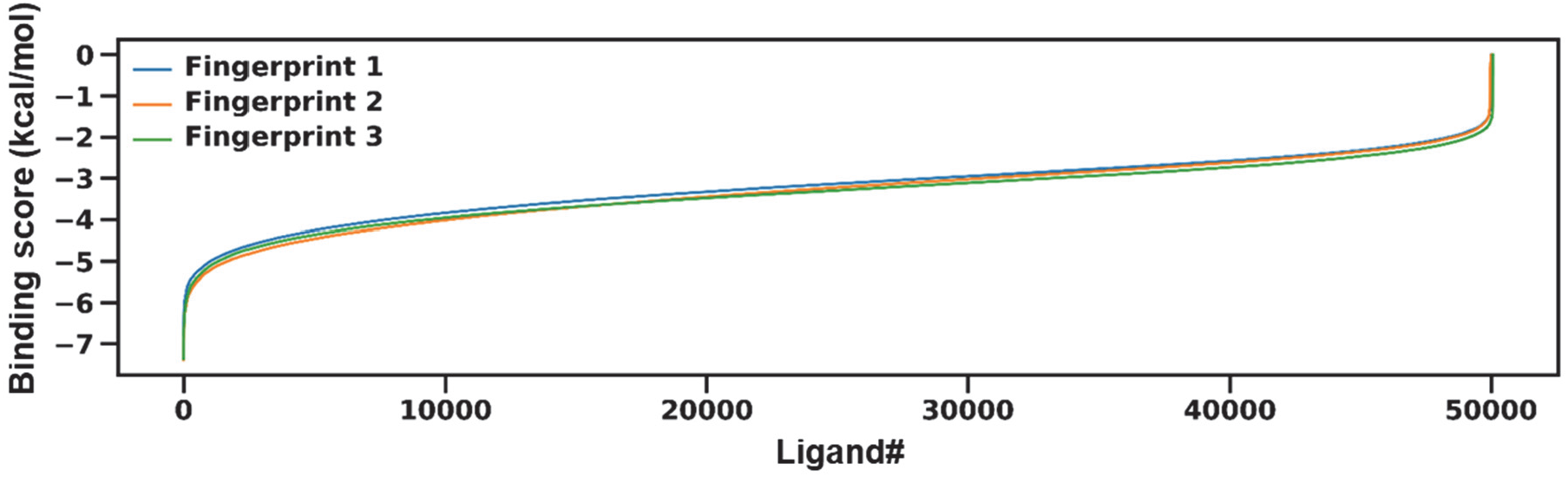
Binding scores of designed drugs. Distribution of binding energies (kcal mol^−1^) for three compound libraries derived from distinct fingerprints. All compounds exhibit favorable binding, with scores below 0 kcal/mol. A small subset of compounds lies near 0 kcal/mol or extends to more favorable negative energies. The three libraries display similar distributions of binding affinities.

### Binding pockets on the MMP1 surface are not at the collagen-specific allosteric residue R405

We identified potential binding pockets with solvent-exposed regions on the MMP1 surface using Fpocket analysis [24]. To this end, we considered whether alpha carbon atoms were solvent-exposed, ensuring that pocket identification was confined to regions physically accessible to ligands. Collagen-bound MMP1 yielded more detected pockets than free MMP1. More pockets for collagen-bound MMP1 are likely due to the presence of the collagen, which contributes additional surface features and interfacial geometries that Fpocket recognizes as candidate binding regions. The statistical distribution of Fpocket’s Drug Scores further differentiates the two systems. Collagen-bound MMP1 exhibits a higher Drug Score of 0.07 ± 0.20 (mean ± SD) across 107 identified pockets. In contrast, free MMP1 displays a lower score of 0.02 ± 0.07 (mean ± SD) across 90 identified pockets.

The highest-scoring pocket for free MMP1, with a Drug Score of 0.48, is near the active site in the catalytic domain, indicating that this region remains the most favorable binding site in the absence of a substrate (**Figure 5A**). In the collagen-bound system, the maximum Drug Score increases to 0.93, indicating the presence of a highly druggable pocket, a larger structural region that extends from the linker region to near the catalytic domain (**Figure 5B**). The minimum Drug Score in both systems is 0.00, with multiple instances observed in each case, indicating that a subset of the detected pockets is not predicted to be druggable under the scoring scheme used by Fpocket. **Figure 5** does not show these low-scoring pockets, although we used all identified pockets as targets, irrespective of Drug Scores, for subsequent drug screening. This approach ensures that the full spectrum of surface-accessible regions is investigated, allowing downstream analyses to evaluate binding behavior across both high-scoring and low-scoring pockets in a consistent and unbiased manner. Note that there are no high-scoring binding pockets near R405 (**Figure 5**), which would normally indicate that R405 is not a druggable site. Nevertheless, as shown next, some drugs designed against dynamic allosteric fingerprints around R405 bind more strongly to R405 than to high-scoring binding pockets in **Figure 5**.

**Figure 5.**
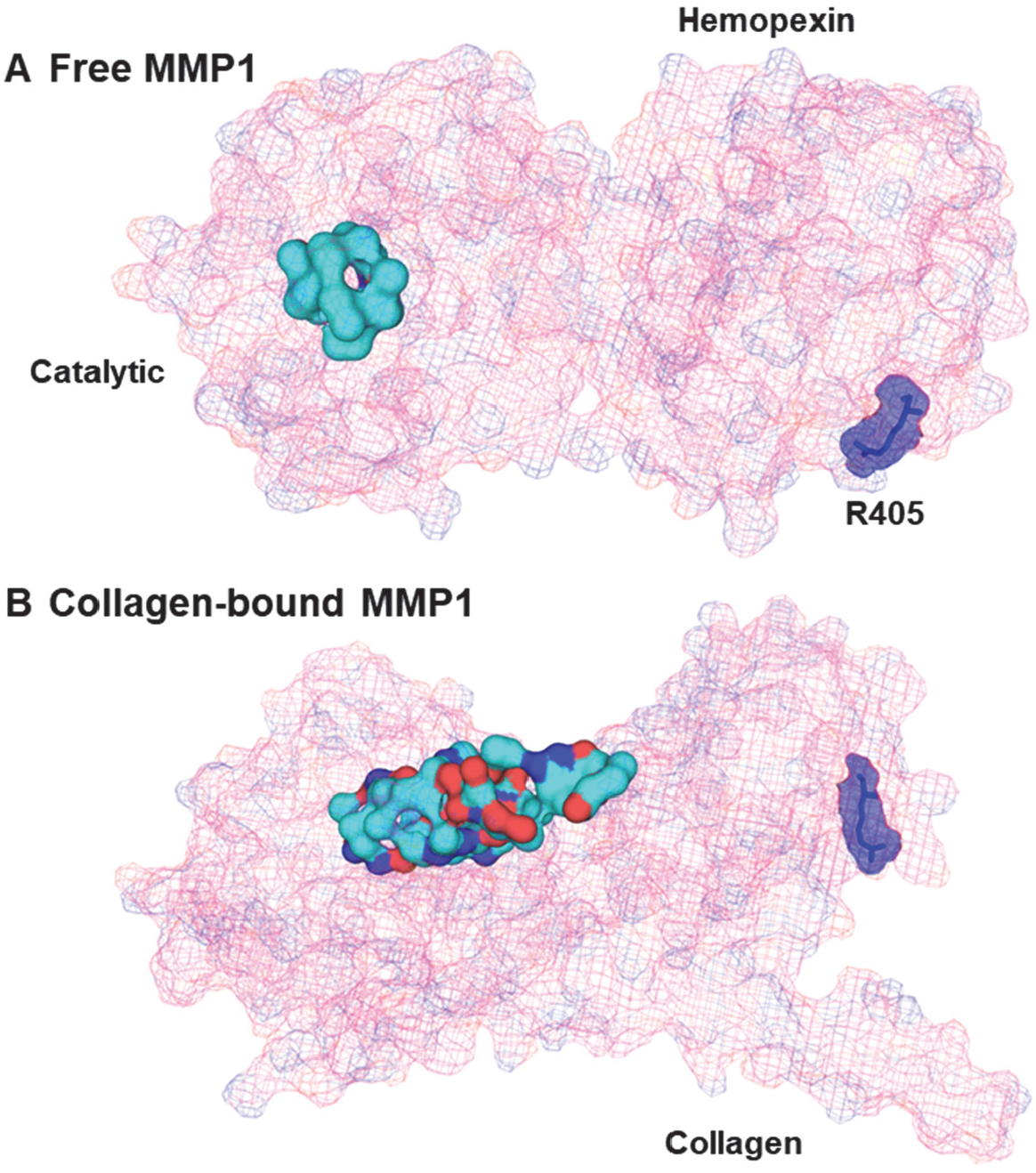
Binding pockets on MMP1. Locations of binding pockets for (**A**) free MMP1 and (**B**) collagen-bound MMP1. In both panels, the full protein is shown in wireframe, with maximal binding pockets rendered as surface representations. The position of R405, corresponding to the allosteric fingerprint site, is highlighted as a dark blue semi-transparent surface.

### Fingerprint-conditioned ligands display preferential binding to the R405 allosteric region compared to other sites on MMP1

We generated 50,000 ligands per fingerprint (**Figure 4**) and selected the top 5,000 based on binding scores. This reduction left three curated libraries of high-affinity, fingerprint-conditioned ligands. For the three fingerprints shown in **Figure 3**, we had a total of 15,000 designed ligands. To quantify the binding affinity of each ligand at R405 and across all identified pockets, we used AutoDock GNINA docking [23] within an area with a 12 Å radius centered on R405. The docking search space was centered on the geometric center of each pocket, and the number of poses per docking run increased to twenty. For each ligand–pocket pair, a representative pose was determined using the median medoid structure. Mean binding affinities were calculated across all twenty poses for each pair. These values were compared to the corresponding affinity obtained at the R405-centered fingerprint for the same ligand. Ligands were retained when their mean binding affinity at the R405-centered site was more favorable than at any alternative pocket, thereby identifying compounds with preferential engagement of the allosteric region at R405 defined by the collagen-bound state.

**Figure 6** shows that the top five ligands bind preferentially near R405 by at least 0.3 kcal/mol relative to competing binding pockets on the MMP1 surface. At body temperature (310 K), this energetic preference corresponds to an estimated ∼1.6-fold longer relative residence time under Arrhenius and Eyring assumptions with comparable prefactors. Although docking scores give a rough estimate of binding energy rather than dissociation kinetics, binding scores represent a lower limit of the energetic barrier that a ligand must overcome for dissociation. These results therefore suggest that preferential binding near R405 may contribute to enhanced kinetic stability and selective occupancy of the collagen-specific allosteric site.

**Figure 6.**
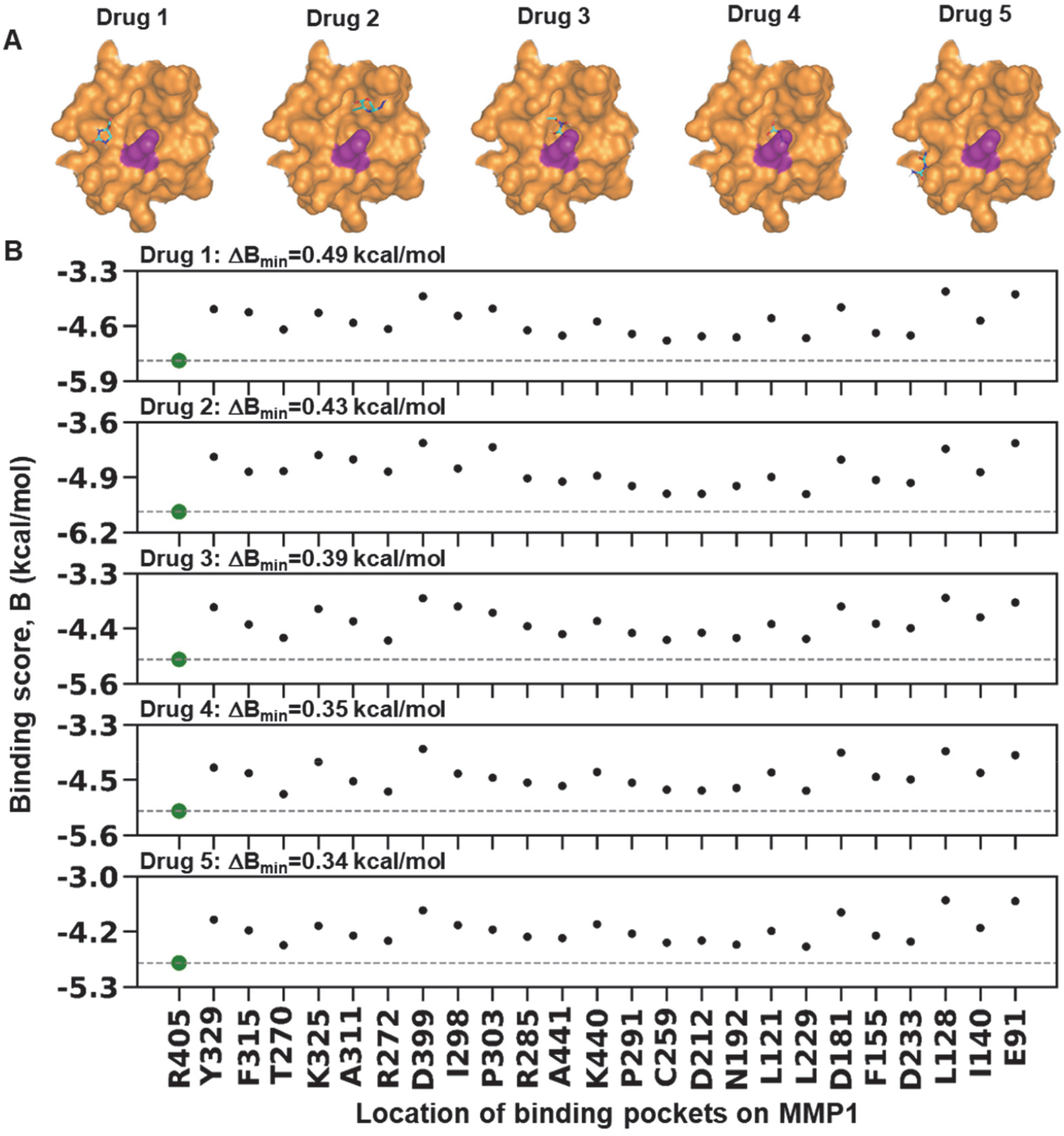
Identification of designed drugs with preferential binding to the targeted site. (**A**) Top five drugs (green sticks) bound to MMP1 near R405 (purple). (**B**) Binding scores of the top five drugs at R405 (green solid circles) and other binding pockets across the MMP1 surface. These drugs bind to R405 with at least 0.3 kcal/mol greater affinity.

## Conclusion

This study establishes a general strategy for preferentially targeting specific protein sites within a broader substrate-specific allosteric drug-design framework (**Figure 7**). We have integrated experimentally validated molecular dynamics simulations, substrate-specific dynamic allosteric fingerprint identification, machine-learning-based ligand generation, and surface-wide selectivity screening to identify ligands that preferentially target specific sites on protein surfaces. Using collagen-bound MMP1 as a model, we identified the top three dynamic allosteric fingerprints near R405 of collagen-bound MMP1 that were structurally distinct from those of free MMP1. These findings support the idea that substrate binding can create functional allosteric states that are not apparent from the unbound enzyme alone.

**Figure 7.**
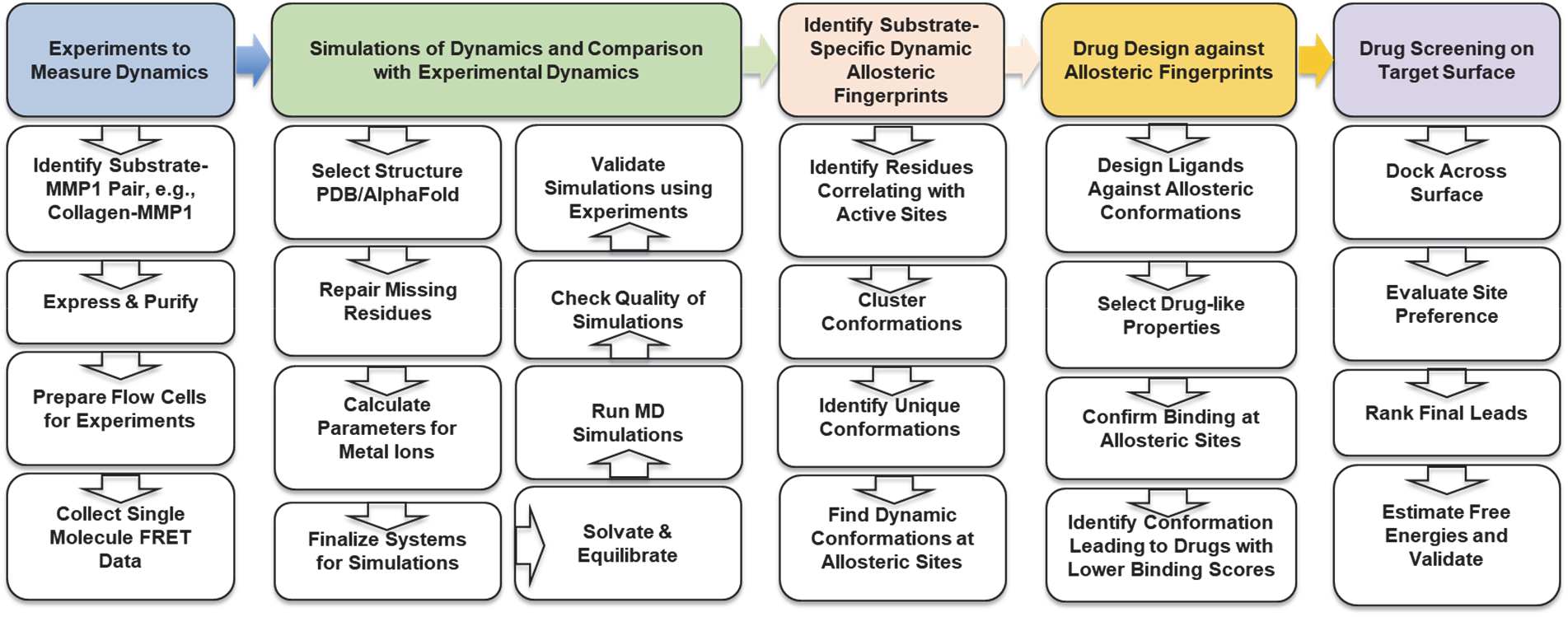
A framework for designing drugs to preferentially target a specific site on MMP1. Two-way feedback between experiments and simulations enables simulation validation. We used validated simulations to identify substrate-specific allosteric fingerprints and to design drugs based on dynamic conformational arrangements at the targeted site. We identified designed drugs that bind more strongly at the target site than at other sites on the protein surface.

We then used these collagen-specific fingerprints to generate ligand libraries and screened candidate molecules not only against the R405-centered site but also across alternative pockets on the MMP1 surface. This surface-wide comparison is important because earlier MMP inhibitor programs were limited by poor selectivity and musculoskeletal toxicity, motivating approaches that avoid broad catalytic-site inhibition [25]. Our results show that fingerprint-conditioned ligands can preferentially bind the collagen-specific R405 region over competing pockets, providing a quantitative route to site-selective allosteric targeting.

Overall, this work addresses a central limitation of conventional *in silico* structure-based drug design: the tendency to optimize ligands against static, predefined pockets without testing whether the same ligands bind more strongly elsewhere on the protein. By combining substrate-dependent conformational selection with full-surface screening, the framework provides a general strategy for designing ligands that target specific sites on proteins. Such substrate-specific allosteric drugs may enable selective modulation of one protein function without altering others, potentially reducing off-target effects and improving precision therapeutic design.

## Materials and methods

### Structural preparation of free and collagen-bound MMP1 systems for simulations

We prepared two systems for simulation: a collagen-bound MMP1 system and a free MMP1 system in which the collagen substrate was removed. Both systems were modeled in the catalytically active form (E219). The free system was generated by removing the collagen from the complex and subsequently equilibrating the protein to remove structural bias associated with collagen binding. We obtained the starting structure from PDB ID 4AUO [3]. We identified missing atoms and residues from the PDB metadata and reconstructed these using Modeller [26]. We generated the catalytically active MMP1 model by modifying the inactive crystal structure by converting a key residue from alanine (A) to glutamic acid (E). No prodomain was included. We capped the termini of MMP1 using ACE at the N-terminus and NME at the C-terminus.

We constructed the collagen model as a periodic molecule spanning the simulation box by forming covalent bonds between the termini of each individual polypeptide chain, such that the molecule crosses one boundary of the unit cell and continues through the opposite boundary. This configuration was used to approximate the rigidity of a collagen fibril, rather than allowing short collagen-like molecules to diffuse in the solvent while bound to MMP1. The collagen triple helix was built to undergo sufficient turns to maintain collagen-like structural behavior, and the system was configured such that MMP1 did not interact with its periodic images. All systems were placed in cubic simulation boxes. For both the collagen-bound and free systems, box dimensions were 7.35 nm × 9.65 nm × 8.93 nm. Periodic boundary conditions were applied in three dimensions. We solvated all systems using the TIP3P water model [27] and added chloride ions to neutralize the system charge.

We described the protein using the AMBER99SB-ILDN force field [28]. We parameterized the catalytic and structural Zn^2+^ and Cu^2+^ ions using values derived from prior quantum-chemical calculations, including force constants, equilibrium bond lengths and angles, and torsional parameters for coordinating ligands.

### MD simulation setup

We performed molecular dynamics simulations using OpenMM (version 8.2.0) [29]. Simulations were executed on the CUDA platform using mixed precision. We maintained the temperature at 295.15 K and the pressure at 1 bar using a Monte Carlo barostat with an update frequency of 25 steps, corresponding to isotropic pressure coupling. We used a Langevin integrator [30] with a time step of 2 fs and a friction coefficient of 1 ps^−1^. Initial velocities were assigned at 295.15 K. We treated long-range electrostatics using the particle-mesh Ewald method with a 0.8 nm cutoff [31]. Short-range nonbonded interactions were truncated at the same cutoff distance. All bonds involving hydrogen atoms were constrained using the HBonds constraint scheme. Periodic boundary conditions were applied to all force terms within the system. We applied harmonic positional restraints to collagen alpha carbon atoms using their initial coordinates as reference positions, with a force constant of 1000 kJ mol^−1^ nm^−2^. Positional restraints were implemented using CustomExternalForce within OpenMM.

### Energy minimization, equilibration, and production simulations

We performed energy minimization using OpenMM’s minimizeEnergy() routine prior to dynamical propagation. We then carried out pre-equilibration to relax structural distortions and stabilize the system. Systems were heated from 0 K to 295.15 K over 1.0 × 10^6^ time steps of 2 fs, corresponding to 100 ps of equilibration. From the minimized structure, three independent simulations were initiated through separate heating procedures, each followed by 40 ns of molecular dynamics equilibration to allow relaxation of the system, including adjustment following collagen removal and equilibration of the unit cell under the applied pressure. Each simulation was then propagated for 500 ns, yielding three independent trajectories per system and a total sampling time of 1.5 μs. Coordinates were written in DCD format using the OpenMM DCDReporter at intervals of 50,000 steps (100 ps). Energies and simulation statistics were recorded at regular intervals during the simulation. Simulation checkpoints were written periodically to enable restart and reproducibility.

### Quality check of simulated conformational dynamics

We analyzed the final 100 ns of each trajectory. We calculated the root mean squared deviation using alpha carbon atoms as the reference selection across this interval for all repeats of the collagen-bound and free systems. The initial frame of each trajectory was used as the reference structure. We calculated the solvent accessible surface area using FreeSASA [32]. SASA was evaluated across the entire structure for all frames in the final 100 ns of each trajectory. Hydrogens were included in the calculation. We used a probe radius of 1.40 Å and applied the default Shrake– Rupley algorithm implemented in FreeSASA.

### Classification of MMP1 conformational dynamics based on solvent-accessibility of R405

We classified frames based on the solvent-accessible surface area near R405. Frames with a surface area less than 5 Å^2^ were classified as buried and excluded. Frames with a surface area between 5 and 30 Å^2^ were classified as partially exposed and excluded. Frames with a surface area greater than or equal to 30 Å^2^ were classified as fully exposed and retained. These thresholds were chosen to distinguish buried, partially exposed, and solvent-accessible states. We used retained frames to construct structurally consistent snapshot ensembles across all trajectories and systems. For each retained frame, we recorded residue identity, chain designation, residue index, and alpha carbon Cartesian coordinates for residues that met the surface-exposure criteria. We calculated the fraction of solvent-accessible alpha carbon atoms per frame and collated frame-wise lists of surface-exposed residues to quantify the prevalence and spatial distribution of solvent exposure across systems.

### Identification of dynamic allosteric fingerprints

We identified candidate binding site fingerprints from the subset of trajectory frames in which R405 was fully solvent-exposed. We generated clustered ensembles of these frames and extracted representative structures. We performed hierarchical clustering using alpha carbon-based RMSD as the distance metric, following rigid-body alignment of the structures using alpha carbons. For each system, we retained the three dominant clusters and extracted representative structures from each cluster. We defined local binding sites by selecting residues within a radius of 12 Å centered on the alpha carbon atom of R405. We retained clusters containing at least 20 frames and excluded smaller clusters. For each retained cluster, we extracted a representative structure and isolated the local binding site. These extracted sites were stored as PDB and SDF files.

### Identification of binding pockets using DeeplyTough

We obtained representative structures by clustering molecular dynamics trajectories for both the free and collagen-bound systems. For each system, we retained three dominant clusters, yielding three candidate conformations per system centered on R405. We compared these binding pockets using DeeplyTough [19]. We performed all-against-all comparisons between fingerprints to generate similarity matrices [33]. We ranked fingerprints by mean similarity score and selected the three lowest-ranking fingerprints per system.

### Ligand generation based on collagen-specific allosteric fingerprints

We performed automated ligand generation using a diffusion-based structure-based design workflow implemented in the GCDM-SBDD framework [18]. Ligand generation was performed only for the collagen-bound system, using the three identified fingerprints. For each fingerprint, we identified binding-site residues by combining solvent-accessibility and spatial-proximity criteria with R405 as the reference. Residues within 12 Å of the reference residue and exhibiting solvent accessibility were retained and used to define the binding environment.

We generated ligands iteratively, producing 500 samples per iteration with incrementally random seeds, and targeted 50,000 unique ligands per fingerprint. We enforced uniqueness using global SMILES-based deduplication. We filtered generated ligands using RDKit-based criteria to ensure chemical validity and removed duplicates. We evaluated ligands using PoseBusters [22], retaining molecules that satisfied at least 80% of validation checks assessing chemical validity, structural integrity, protein–ligand steric compatibility, geometric strain, and topological consistency. Ligands passing all filters were aggregated into a master dataset for each fingerprint.

## Acknowledgments

This work was supported by a grant to S.K.S. from the National Institutes of Health (GM145210).

## Author contributions

S.K.S. conceived and designed the overall project. A.N. conceived an approach to drug design based on molecular dynamics and machine learning. A.N. ran MD simulations, implemented machine learning-based drug design, and screened drugs. A.N., C.H., and S.K.S. analyzed the results. S.K.S., A.N., and C.H. wrote the manuscript. All authors edited the manuscript.

